# PiFlow: A Biocompatible Low-Cost Programmable Dynamic Flow Pumping System Utilizing a Raspberry Pi Zero and Commercial Piezoelectric Pumps

**DOI:** 10.1101/192047

**Authors:** Timothy Kassis, Paola M. Perez, Chloe J. W. Yang, Luis R. Soenksen, David L. Trumper, Linda G. Griffith

## Abstract

With the rise of research utilizing microphysiological systems (MPSs), the need for tools that enable the physiological mimicking of the relevant cellular environment is vital. The limited ability to reproduce crucial features of the microenvironment, such as surrounding fluid flow and dynamic changes in biochemical stimuli, severely limits the types of experiments that can be carried out. Current equipment to achieve this, such as syringe and peristaltic pumps, is expensive, large, difficult to program and has limited potential for scalability. Here, we present a new pumping platform that is open-source, low-cost, modular, scalable, fully-programmable and easy to assemble that can be incorporated into cell culture systems to better recapitulate physiological environments. By controlling two commercially available piezoelectric pumps using a Raspberry Pi Zero microcontroller, the system is capable of producing arbitrary dynamic flow profiles with reliable flow rates ranging from 1 to 3,000 µL/min as specified by an easily programmable Python-based script. We validated the accuracy of the flow rates, the use of time-varying profiles, and the practicality of the system by creating repeatable dynamic concentration profiles using a 3D-printed static micromixer.

## Specifications Table

|  |  |
| --- | --- |
| Hardware name | *PiFlow* |
| Subject area | Biological Sciences |
| Hardware type | Biological sample handling and preparation |
| Open Source License | *Attribution-ShareAlike 4.0 International (CC BY-SA 4.0)* |
| Cost of Hardware | \$350 |
| Source File Repository | https://osf.io/fdejw |

## 1. Hardware in Context

With the rise of organs-on-chips (OOC) and microphysiological systems (MPS)[1,2,11,3–10], the ability to mimic the highly dynamic cellular microenvironment has become crucial in allowing these systems to recapitulate in vivo physiological responses[12]. Two critical factors involved in mimicking the cellular microenvironment are dynamic flow rates and variable concentration profiles of hormones, proteins, sugars and other biochemical stimuli[13]. For example, mechanical stimuli, such as flow-induced wall shear stress[14,15], and biochemical stimuli, such as varying hormone levels[16], have been shown to dramatically improve the functionality of these in vitro model systems and their physiological relevancy. While there are numerous techniques to impose flow in microfluidic setups[17], existing methods face varying limitations. Continuously imposing adequate flow for a long period of time requires large and complex setups that involve external syringe pumps, peristaltic pumps or the use of one of the commercialized microfluidic pumps. While each of these configurations offers certain advantages, they are typically expensive, tedious to assemble and setup, allow for limited programmability, and do not generally lend themselves as scalable solutions due to their large footprint and excessive cost. While our proposed system (referred to as PiFlow) is still far from ideal, we believe it will be useful in a variety of contexts and will provide several advantages for a diverse set of laboratory applications.

## 2. Hardware Description

There have been numerous attempts made recently by academics to address some of the limitations of current pumping systems[17,18,27,28,19–26]. Examples include an open-source syringe pump[20], a low-cost syringe pump[21], and a scalable pumping system using a Braille display as an actuator[29,30]. Here we present a microfluidic pumping platform designed to offer greater ease of assembly, higher scalability and customized flow-rate programmability compared to other currently available commercial and academic systems. Our modular sub-$350 setup incorporates two commercially available piezoelectric pumps[31] capable of imposing reliable flow rates ranging from 1 to 3,000 µL/min. The pumps are controlled via a Wi-Fi enabled Raspberry Pi Zero microcontroller and can be accessed remotely while in the cell culture incubator using a remote server. The entire system has a minimal footprint, and the 3D-printed casing is designed to make the individual devices stackable. Because of the low power requirements, the system can be battery operated, thus allowing for mobility and uninterrupted flow during day-to-day use as well as eliminating the need for external connections to the incubator (such as pneumatic lines, power cables and fluid tubing). We demonstrated the system’s utility by integrating it with a custom-designed 3D-printed microfluidic static mixer and by programmatically controlling the flow-rates on the two pumps to create dynamic concentration profiles that can potentially be used to impose time-varying concentrations of hormones, drugs or other biochemical stimuli in the designated OOC or MPS on time scales ranging from seconds to weeks.

We designed PiFlow to address many of the limitations that we ourselves encounter every day when working with external pumping systems for microfluidic devices. While commercially-available flow platforms can offer numerous benefits, systems such as syringe and peristaltic pumps have always been cumbersome to use, have a large footprint, thus making them difficult to scale, are difficult if not impossible to program, and are costly for biological experiments that require a high number of replicates and sample conditions. To address these limitations we set out to design a pumping platform specifically geared towards flexibility, programmability, and improved scalability. The result is a modular pumping system where each module is composed of two piezoelectric pumps, two driving circuits, a custom designed PCB and a 3D printed casing (**Figure 1A**). Having two programmable pumps on each platform greatly expands the utility of the system as it allows for applications involving multiple flow rates such microphysiological systems with apical and basal flow, and two-solute mixing applications. Each module is controlled using a single Raspberry Pi Zero W computer that offers a full-fledged Linux operating system and Wi-Fi capabilities. This permitted us to remotely access the PiFlow module using a VNC server free of charge (see the official Raspberry Pi documentation for using RealVNC.com) while the system is in the incubator, thus allowing both real-time monitoring and full programming capabilities using a number of languages such as Python, Scratch, JAVA, C and C++. PiFlow allows the user to create Python scripts that control flow rates for each pump individually. This capability can greatly reduce user intervention in biological experiments by potentially automating many fluid addition and exchange tasks. The pump control scripts that we developed for the module were all in Python and are available with an open source license[32,33]. Each module has a small footprint (122x42x39 mm) that makes it easy to handle and to place in a cell culture incubator with no external wiring or tubing if used with an external USB power-pack (**Figure 1B**). We demonstrated that the entire electronic assembly is robust to the high humidity found in a typical cell incubator by running the platforms continuously for 10 days without any perceived side effects to the hardware or reduction in performance (**Figure S9**). The Bartels Microtechnik mp6 piezoelectric pumps were chosen due to their small size, reliability, high dynamic range, low power consumption (<200 mW), long operating life-time (~5,000 hrs) and chemical compatibility. Each pump has a single barbed inlet and outlet (OD = 1.9 mm) making it easy to interface to using readily available standard commercial tubing[34]. Additionally, the pump material that is in fluid contact is biocompatible[34] where the inner surface of the pump is made from polyphenylene sulphone (PSSU) and is both sterilizable using 70% ethanol and autoclavable. The driving circuits generate a 270 Vpp voltage from the 3.3 V supply voltage of the microcontroller and can be used to control both the amplitude and frequency of actuation. We found that controlling just the frequency was sufficient. The frequency was set using a software pulse-width modulated (PWM) signal with a duty cycle of 95% generated by two output pins of the microcontroller and supplied to each of the CLOCK pins of the mp6-OEM. We designed the casing with a sliding mechanism that allows the user to stack multiple modules on top of each other depending on how many total pumps are required (**Figure 1C**). The end result is a low-cost (<$350), fully programmable, remotely-accessible, battery-operated, scalable and easy-to-operate pumping platform that can be utilized for a variety of fluidic applications. Given the biocompatible nature of PiFlow, it opens up the possibility to automate a variety of biological protocols.

**Figure 1:**
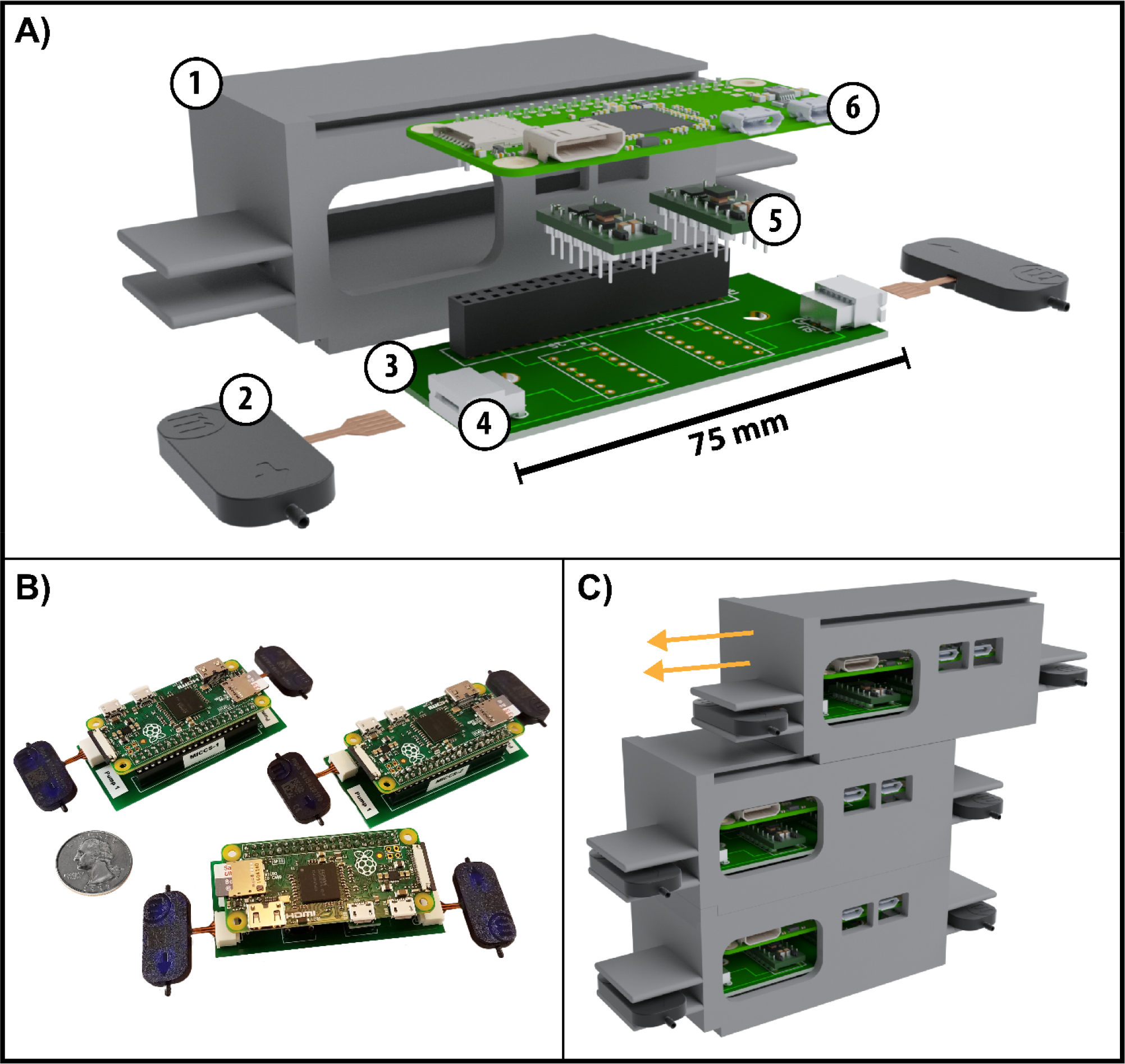
Overview of the PiFlow system. **A)** An exploded rendering of the complete module showing a (1) 3D printed case, (2) the commercially available mp6 piezoelectric pumps, (3) the custom designed PCB, (4) pump connectors, (5) commercially available driving circuits and (6) the Wi-Fi enabled Raspberry Pi Zero Linux computer. **B)** A US quarter next to three modules to show the small footprint. Each module can be given a host name and accessed remotely via a free remote server using RealVNC.com. **C)** The casing allows the modules to be easily stacked when more than two pumps are required. Given the cost, ease of assembly and size of the footprint of PiFlow, it is possible to fit several dozen modules inside a typical cell culture incubator.

## 3. Design Files

| *Design file name* | *File type* | *Open source license* | *Location of the file* |
| --- | --- | --- | --- |
| PCB | PCB CAD | CC BY-SA 4.0 | https://osf.io/p5gfb |
| Casing | STL | CC BY-SA 4.0 | https://osf.io/gyuwk |
| Mixer Mold | STL | CC BY-SA 4.0 | https://osf.io/5a8ur |
| Battery Casing (optional) | STL | CC BY-SA 4.0 | https://osf.io/b8tev |
| CFD Mixer Simulation | COMSOL Multiphysics | CC BY-SA 4.0 | https://osf.io/q2rxm |
| Supplementary Figures | PDF | CC BY-SA 4.0 | https://osf.io/s7frp |
| Code | Python | MIT | https://github.com/TKassis/PiFlow |

- ***PCB*** is a CAD file for a ready-to-fabricate printed circuit board. It is a 2-layer board that can be outsourced to any third party fabrication facility of your choice or made in-house if the right equipment is available.
- ***Casing*** is a stereolithography (STL) file for the case that houses all the electronics and pumps (**Figure 1A-1**). It can be printed on any 3D printer.
- ***Mixer Mold*** is a stereolithography (STL) file for the micromixer. The mold can be either micromachined or 3D printed. It is used with PDMS (see BOM) to make the micromixer.
- ***Battery Casing*** is an optional stereolithography (STL) file for the casing for the specific USB power-pack we used. The main advantage is that it slides into the main casing for convenience.
- ***CFD Mixer Simulation*** is a proprietary format, but optional, multi-physics COMSOL file (MPH version 5.3) that allows you to model the dynamic concentration profiles resulting from your programmed flow rates.
- ***Supplementary Figures*** is a PDF file with extra photos of the components and the system in operation as well as experimental validation of robustness to incubator conditions.
- ***Code*** is a set of python scripts used to operate the platform. Available via GitHub with an MIT license.

## 4. Bill of Materials

| *Designator* | *Component* | *Number* | *Cost per unit - USD* | *Total cost- USD* | *Source of materials* | *Material type* |
| --- | --- | --- | --- | --- | --- | --- |
| *Raspberry Pi* | *Raspberry Pi Zero W with header* | 1 | 14 | 14 | Adafruit | *Non-specific* |
| *Piezoelectric pump* | *mp6 - piezoelectric pump* | 2 | 38 | 76 | Servoflo | *Non-specific* |
| *Driving circuit* | *mp6-OEM driving circuit* | 2 | 63 | 126 | Servoflo | *Non-specific* |
| *Check valve (optional)* | *mp6-CV – check valve* | 2 | 50 | 100 | Servoflo | *Non-specific* |
| *PCB* | *Custom component* | 1 | 10 | 10 | PCBWeb | *Non-specific* |
| *PDMS (optional)* | *Sylgard® 184 Silicone Elastomer* | 1 | 0.50 | 0.50 | Corning | *Polymer* |
| *Molex connector* | *FCC Connector, Right-Angle, 4 Circuits (39-53-2044)* | 2 | 1 | 2 | Molex | *Non-specific* |
| *Female header* | *Hammer Header Female - Raspberry Pi Connector* | 1 | 3.25 | 3.25 | Adafruit | *Non-specific* |
| *Casing* | *Custom component* | 1 | 2 | 2 | Printed on a Form 2 | *Polymer* |

Additionally, the following equipment might be needed:

- A 3D printer for printing the casing (can also be outsourced to companies such as Shapeways, i.materialise, Sculpteo or 3D Hubs).
- A soldering iron for soldering the components to the PCB.
- An oven and plasma cleaner for fabricating and bonding the micromixer (optional).

## 5. Build Instructions

### Assembling the Components and Initial Setup

PiFlow is composed of two piezoelectric pumps, two pump driving circuits, two Molex connectors, a printed circuit board (PCB) and a Raspberry Pi Zero microcontroller (**Figure 1**). An optional check-value can be integrated in line with the tubing to limit backflow if needed. Two commercially available mp6-OEM driving circuits were used in conjunction with a Raspberry Pi Zero W running Raspbian Jessie Linux distribution to control both pumps. A custom-built PCB (PCBWeb Designer) along with two 1.25 mm pitch Molex connectors were used to connect all the various components via soldering. To assemble the main PiFlow device:

1. Solder the Molex connectors, female header and driving circuits onto the PCB board (**Figure 2**).
2. Push in the pumps (copper contact side facing up) and push in the Raspberry Pi Zero into the header (**Figure 3**).
3. Set up the Raspbian operating system on the Raspberry Pi Zero using the well documented procedure thoroughly described by the Raspberry Pi Foundation here.
4. Download the PiFlow repository directly to your Raspberry Pi from GitHub.

**Figure 2:**
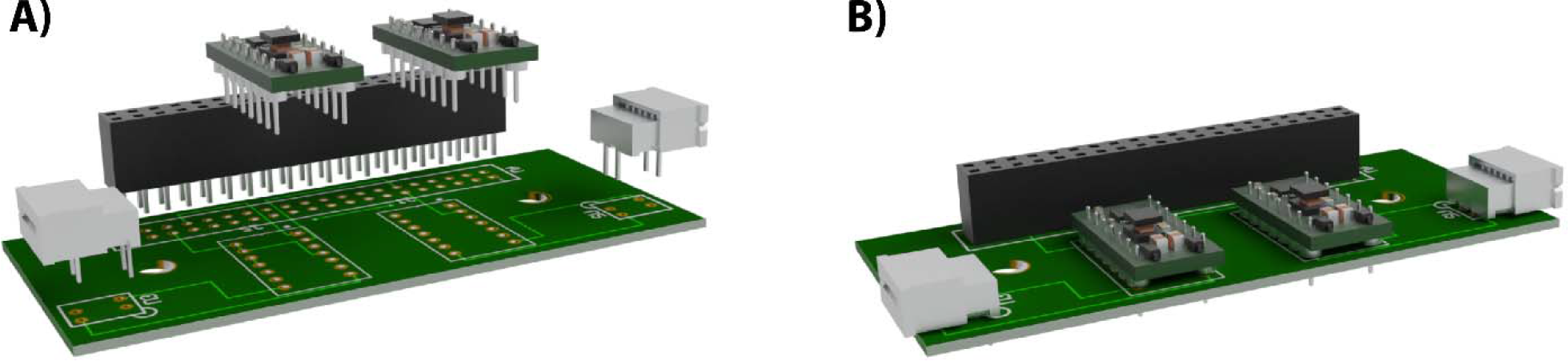
First step of device assembly. **A)** Presoldering. **B)** What the device should look like after soldering the indicated components.

**Figure 3:**
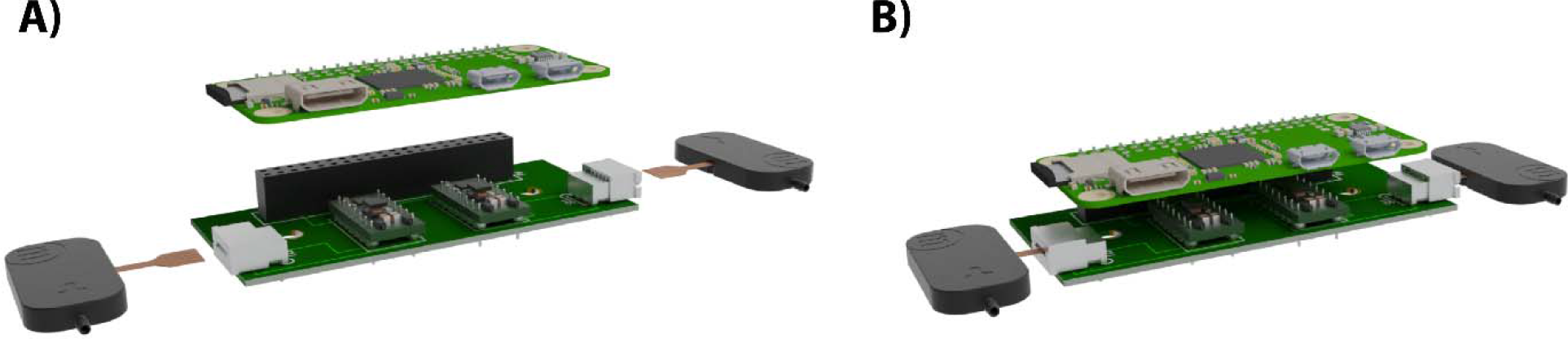
Second step of device assembly. **A)** Once the components are soldered in you can push in the Raspberry Pi (that already has male headers) and the pumps. **B)** The system is now ready for software installation.

The main assembly is now complete and the device can be used (please see operation instructions). Additional more detailed 3^rd^-party instructions are available from the following sources:

- The piezoelectric pumps (cleaning, maintenance, specs…etc)
- Raspberry Pi (set-up, installation, updates…etc)
- GitHub (downloading the code, contributing…etc)
- Soldering tutorial

### 3D Printing the Casing

The casing and the sliding mechanisms were designed using Autodesk Fusion 360 and exported as an STL file (provided) ready for printing on any available 3D printer according to normal manufacturer’s instructions. The following instructions are for a Form 2 3D printer (Formlabs, Somerville, MA):

1. Load the STL casing file into the Formlabs PreForm software and print at a vertical resolution of 50 µm using any Formlabs standard resin (we have tried both ‘gray’ and ‘clear’).
2. When the print is complete soak in Isopropyl Alcohol (IPA) for 20 minutes, then dry with an air gun.
3. Remove the support structures.
4. Post-cure under UV light for 1 hour using a Salon Edge 36W Professional UV dryer (Amazon) or any other available UV source. Alternatively, leave under sunlight for several hours.

PiFlow can now be placed inside the casing. Note that the pumps have to be removed first then put in after insertion into the casing.

### Fabricating the PDMS Micromixer (Optional)

The mixer was made from poly(dimethylsiloxane) (PDMS) bound to a glass slide. While PDMS is a highly absorptive material, we chose to use it to maintain a low cost of fabrication and operation of PiFlow. The mold was designed using Autodesk Fusion 360, exported as an STL file and printed with standard clear resin on a Form 2 3D printer at a set vertical resolution of 25 µm. The print was then soaked in IPA for 20 minutes, washed thoroughly with fresh IPA, and then dried with an air gun. After removing the support structures, the mold was post-cured under UV light for 1 hour using a Salon Edge 36W Professional UV dryer. To construct a device,

1. Thoroughly mix PDMS at a 10:1 ratio of base to curing agent (Sylgard 184, Dow Corning) and pour into the 3D printed mold.
2. Degas for 20 minutes vacuum desiccator.
3. Cure in an oven at 60 °C for a minimum of 8 hours.
4. Note: The first time a new 3D-printed mold is used, PDMS does not fully cure due to resin leaching out of the 3D print and should therefore be discarded.
5. Punch inlet and outlet holes for tubing using a 1 mm biopsy punch.
6. Clean the PDMS device with scotch tape.
7. Air plasma treat for 30-60 seconds and immediately bond to a 1 mm thick glass slide.
8. Leave in oven at 60 °C for minimum of 2 hours to strengthen the PDMS-glass bonds.
9. If to be used for cell culture applications, sterilize by soaking in 70% ethanol for 5 minutes, then rinsing thoroughly with sterile water.
10. Optionally, before use, mixers should be filled with 1% bovine serum albumin (BSA) solution for 1 hour to decrease non-specific binding.

The micromixer can now be used for mixing applications if needed.

## 6. Operation Instructions

Python 3.5 along with the Raspberry Pi GPIO library were used to control a software PWM signal generated through the Raspberry Pi Zero W. Pins 16 and 19 were used as power pins and pins 17 and 18 as output pins for the PWM signal (see PCB file). The frequency of the PWM was modulated to control the flow rate. Two different output pins were used to generate a PWM signal for each pump. A calibration curve (frequency vs. flow-rate) was created to determine the PWM signal corresponding to each flow rate. A small Python library was developed with two main functions: *Constant_Flow(Q1,Q2,T)* generates a constant flow rate (Q1 and Q2 for pumps 1 and 2 respectively) for a specified duration (T in sec), *and Dynamic_Flow(csv_path)* takes in a file path for a comma separated “.csv” file. The “.csv” file can be generated by the user in any way they see fit to perform any custom pump functionality by dynamically controlling both pumps. All Python code used in this paper is available at *https://github.com/TKassis/PiFlow* and can be downloaded directly to the Raspberry Pi.

### 6.1. Basic Pump Flow Usage

Follow the following steps to get up and running with PiFlow:

1. Connect to the Raspberry Pi and open the *PiFlow_Program.py* file.
2. To operate PiFlow at constant flow rates, simply input the flow rates in µL/min for the two pumps respectively along with the total duration to run in seconds. For example, to run pump 1 at 200 µL/min and pump 2 at 600 µL/min for 6 hours (21,600 seconds) the *PiFlow_Program.py* file would look like the following:

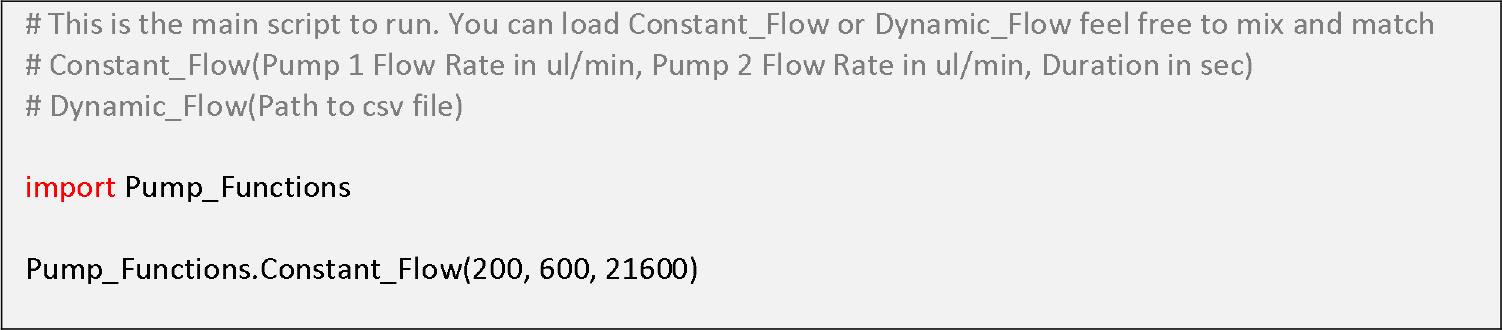
3. Run the Python script and the pump should start running.

For dynamic discrete programmable flow rates the user can have a sequence of the above functions. For example to create a flow profile where each pump runs for 2 hours at 500 µL/min then stops for 6 hours then runs again at 200 µL/min for 12 hours:

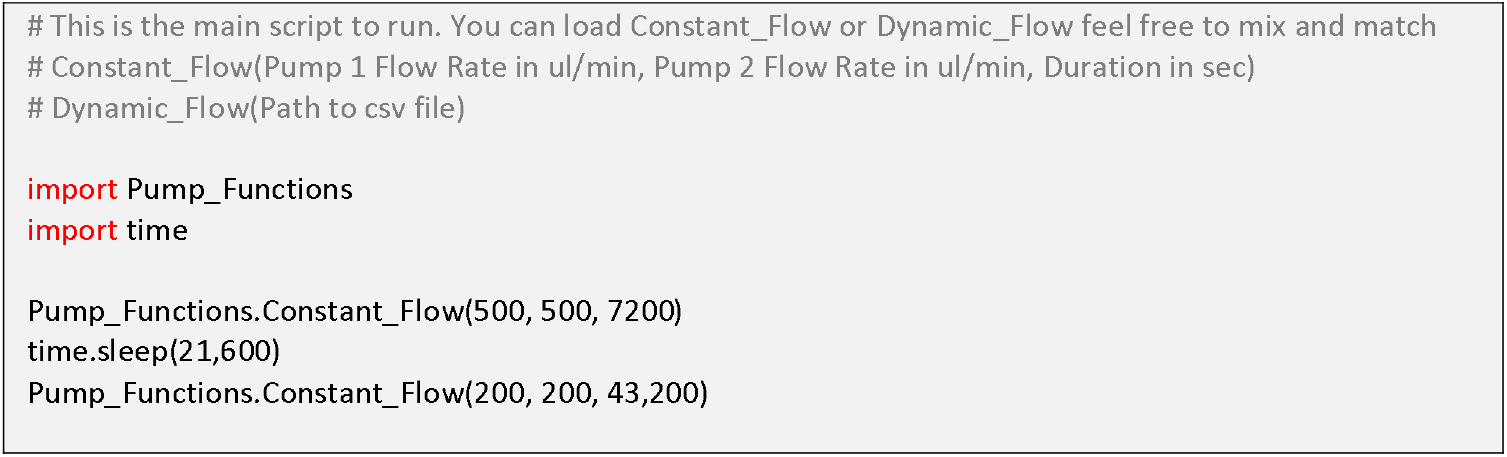

To create a truly dynamic flow profile the user has the option of using a “.csv” file. The csv file should contain three columns with the sampling interval in seconds in the first column and the flow rates, in µL/min for each pump, in the next two columns (please see example “.csv” file in supplemental). An example code would be:

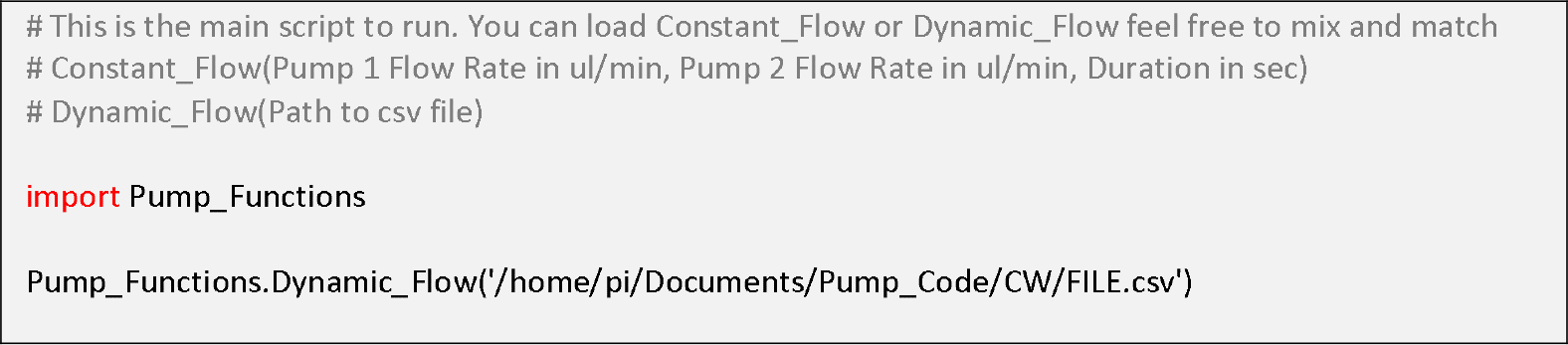

## 7. Validation and Characterization

### Calibration Offers Reliable and Reproducible Flow Rates with a High Dynamic Range

The pump flow-rate is set by specifying a PWM signal with a selected frequency. A calibration curve was created by weighing the total pumped volume, given a certain frequency input, of ultra-filtered water for selected time intervals. Each measurement was repeated three times. A calibration curve was created for each pump separately. The flow rates were then validated under the same conditions in which the subsequent experiments would take place. For the flow validation experiments, the mass of water was measured for 10-minute flow intervals and measurements were repeated four times for each prescribed flow rate. The two pumps on each PiFlow module were calibrated separately. While the two calibration curves showed the same linear trend, there is a small gain difference between them which requires each pump to have its own calibration curve. This gain difference is most likely due to manufacturing variability of the pump geometry. The calibration data for both pumps is linear up to 100 Hz, at which point there is an abrupt change in slope. The cause of this change in slope is undetermined, but is consistent with the manufacturer’s documentation. To account for the change in slopes, we divided the curve into two sections. We fit a straight line through each set of data points where the first line was from 0-100 Hz and the second line was from 100-300 Hz (**Figure 4A,B**). The calibration values were subsequently used in the pump control code. We validated the resulting code by measuring the flow rate using mass of water collected over a 10-min period for one of the pumps (**Figure 4C**), and we found the flow rate to be highly accurate even at low flow rates (**Figure 4D**). The validated flow rates illustrate the high dynamic range possible with PiFlow, ranging from 1 to 3000 µL/min.

**Figure 4:**
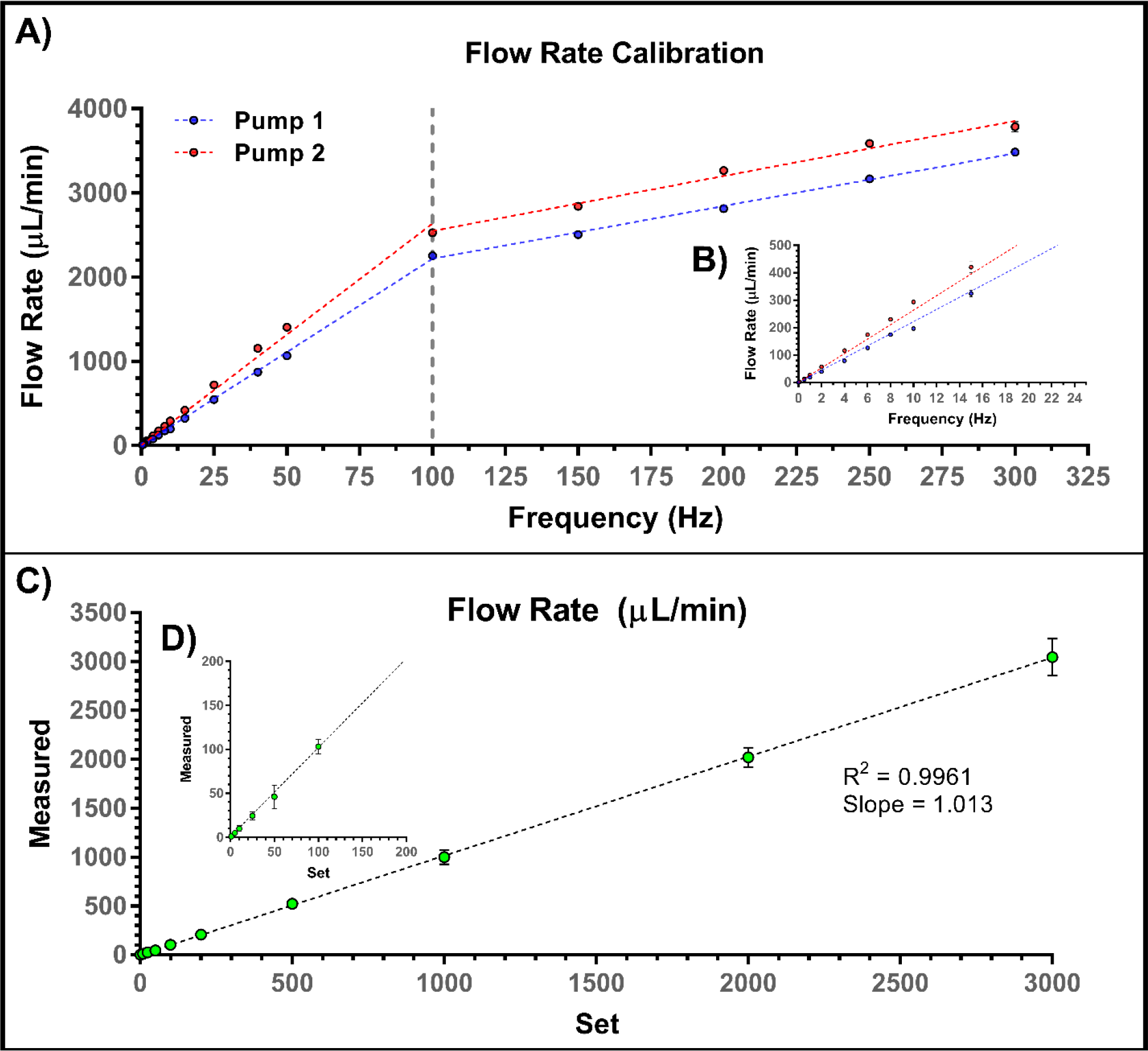
Calibration and Validation of Flow Rates. **A)** Calibration curve for a PiFlow module for each of the two pumps created by measuring the mass of water displaced at given set frequencies (n=3, error bars indicate standard deviation). The response exhibits two linear intervals from 0-100 Hz and from 100-300 Hz. Curve details not provided here as they need to be established for each set-up independently (see code for details). **B)** Close-up of the low flow-rate region **C)** Validating the flow rates of the specific setup used for the subsequent experiments showed reproducible flow rates even down to 1 µL/min **(D)** (n=4, error bars indicate standard deviation).

### Precise Repeatable Dynamic Concentration Profiles Created using a 3D-Printed Mixer and Computational Model

The ability to create dynamic flow waveforms has been crucial in many in vitro physiological studies such as those investigating effects of blood[35–37] and lymphatic shear stress[38,39]. Several systems have recently demonstrated the ability to control flow dynamically or through valving techniques to create temporal concentration profiles[18,40–48]. While each of them is powerful for certain applications, we believe that the flexibility PiFlow provides allows researchers to more easily accomplish such tasks. PiFlow can reliably produce many dynamic flow profiles (<3,000 µL/min peak flow rate) by loading a custom “.csv” file containing the desired flow rates and time intervals. As an example, we chose here to demonstrate the improved functionality of PiFlow by incorporating a microfluidic static mixer to create dynamic concentration profiles useful in many cell stimuli studies[13] whether the stimulus be drugs, glucose[49,50], hormones or growth factors[41,43]. By connecting one pump to a reservoir containing a concentrated solution and the other pump to a reservoir containing buffer solution, dynamically changing the ratio of the flow rate of one pump to the other, and mixing the flow from both pumps, we can dynamically control the concentration of a solute while keeping the flow rate constant (**Figure 5A**). If desired, one can also have a dynamically changing flow rate. While PDMS has many undesirable absorptive properties that make it a bad choice for a mixer (especially when used with steroid hormones or in drug studies) there are many commercially available thermoplastic-based static mixers that can reliably mix two solutions. We chose to design our own, utilizing a 3D printed microfluidic mold[51–53], in order to demonstrate not only the functionality of PiFlow, but also to show the ability of rapidly producing microfluidic devices for use with the system without the need for expensive microfabrication techniques such as photolithography[54] or micromachining[55]. The resulting device was composed of a series of interconnected triangles spanning the two inlets to the single outlet (**Figure 5B**). The expansion of fluid into the triangle region causes the formation of small vortices that assist with mixing (**Figure S7**). To validate the mixing efficiency of the static mixer we conducted 2D multi-physics simulations using COMSOL 5.3. The study was composed of two COMSOL simulation steps. For the first step, we solved a discretized (P2 + P1) laminar flow simulation where the inlet velocity (0.055 m/s) for each of the inlets was specified to correspond to a 400 µL/min flow rate. The second step utilized the laminar flow solution and a discretized quadratic transport of dilute species study to solve for mixing throughout the device given a diffusion coefficient of 4.25x10^-10^ m^2^/s for fluorescein[56]. The concentration of fluorescein was set to 1 mol/m^3^ for the high-concentration inlet while the other inlet had pure water. Before any experimental validation, we simulated the mixing of fluorescein with water at a 400 µL/min steady flow rate for each inlet (**Figure 5C**) and showed that we achieve complete mixing after passing a minimal distance (**Figure 5D**).

**Figure 5:**
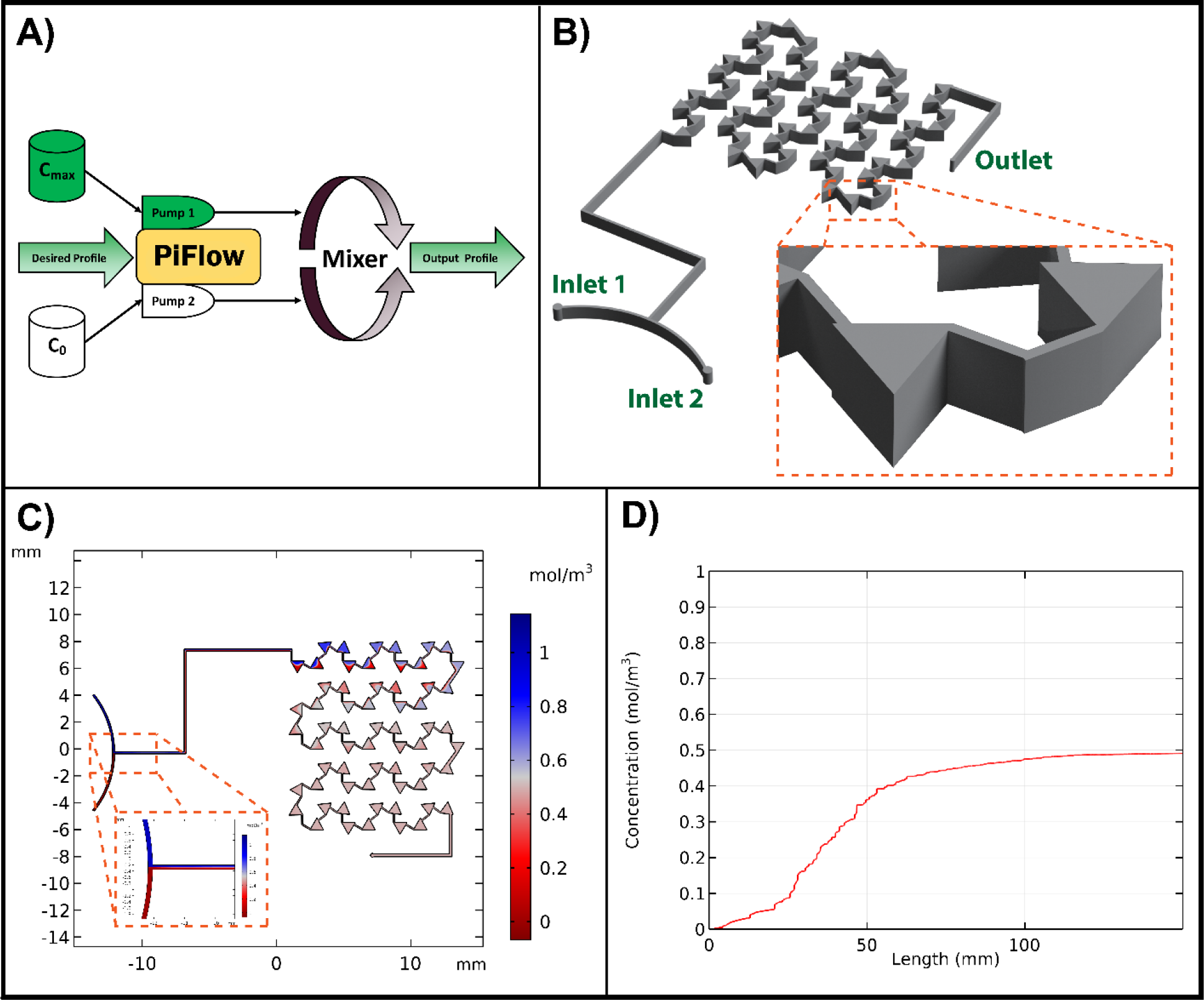
Simulation of Mixer Performance using PiFlow. **A)** By utilizing two reservoirs, each connected to one of the pumps and then to a static efficient mixer, we can create dynamic concentration waveforms while holding the output flow rate constant. **B)** A CAD model of our microfluidic mixer that can handle a large range of input flow rates from a few µL/min to 3,000 µL/min and above. **C)** A CFD simulation showing efficient mixing of fluorescein at 37 °C under 400 µL/min flow rates for each of the inlets. **D)** Full mixing occurs after 125 mm at these given flow rates.

In addition to dynamically perturbing mono-cultures of cells to probe for mechanistic insights, being able to easily prescribe dynamic concentration profiles offers new capabilities for stimulating organ-on-chip and microphysiological systems. By uploading a “.csv” file, we can recreate any arbitrarily defined concentration profile. Utilizing the same computational fluid dynamics (CFD) and solute transport model used for the constant flow above, we performed a parametric sweep over set values of inlet velocities to simulate dynamic concentration profiles. We found that our simulation accurately predicted the concentration profiles given our programmed flow rate values derived from the required concentration profiles (**Figure 6**). We chose to produce three different waveforms (ramp, sinusoid and pharmacokinetic-like) over a 10 minute period for validation. The user, however, can create profiles on any time scale (from seconds to weeks) depending on the volume, stability, and other limiting factors of the solutions to be used. It is important to note that the used pumps are inherently pulsatile in nature with a stroke volume according to the manufacturer of around 2 µL. This pulsatility was not incorporated into the simulation as the phase difference between the two pumps cannot be controlled accurately, thus although the real concentration values at the output will likely be very similar to the simulation, the extent of spatial mixing along the length of the device might not necessarily be reflected accurately by the simulation in **Figure 5D**. The computational model, along with a microfluidic mixer, allows us to quickly validate the resulting output concentration profile for any given solute, temperature, inlet flow rates and duration thus reducing the need to validate these concentration profiles before an experiment through physical measurements of the solute which can be prohibitively costly, tedious and time-consuming.

**Figure 6:**
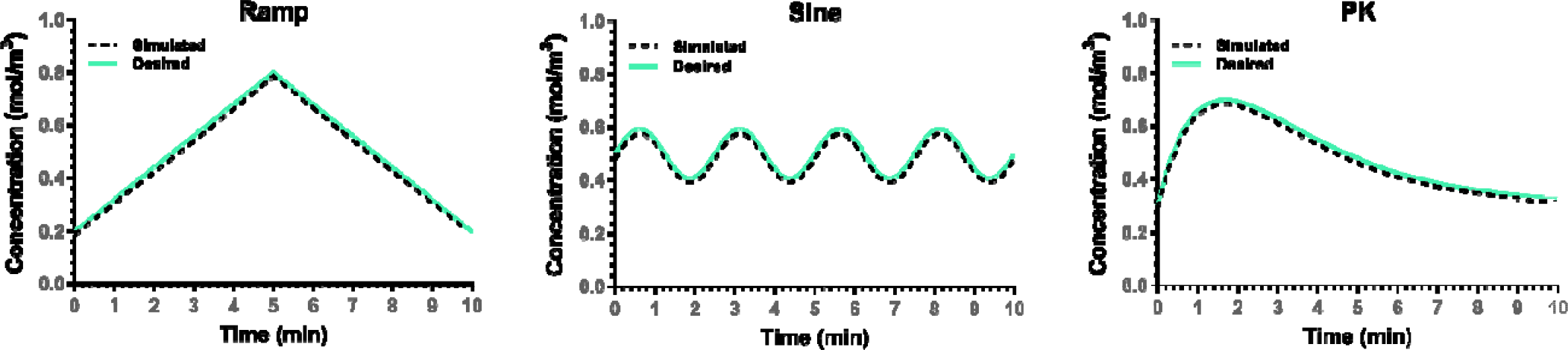
Validation of the flow rate “.csv” file to achieve desired concentration using the CFD model. The developed CFD model allows us to computationally validate that our prescribed flow waveforms create the desired concentration profiles, thus minimizing the need for real measurement-based validation before a stimulation experiment.

To experimentally validate that we were in fact achieving the desired concentrations and that both our hardware and computational model were accurate, two 250 mL bottles were filled with deionized water and deionized water with stock concentration of fluorescein respectively (**Figure 7A**). The setup, using PiFlow connected to the mixer, was placed in a temperature-controlled incubation chamber in a Zeiss LSM 880 confocal microscope. Images were acquired every 5 seconds with a resolution of 512x512 using ZEN Black confocal software (Zeiss) and at a 16-bit depth resolution. The images were saved as .czi files Each file was loaded into the open-source image processing software, Fiji[57,58], and fluorescence intensity was measured using Fiji’s built-in region of interest (ROI) intensity measurement functionality The ROI was drawn as a small rectangle in the center of the outlet channel of the mixer. The intensity measurement that Fiji calculated was the mean intensity of all the pixels within the ROI. Using a calibration curve (mean fluorescence intensity vs. fluorescein concentration) that was created using the same setup for known fluorescein concentrations, the final concentration values were determined from the fluorescence intensity. With the same flow rate values used in the simulations, we achieved the same concentration profiles predicted by the computational model when measuring fluorescence intensity of fluorescein. The concentration profiles were highly accurate and reproducible with an R^2^ > 0.97 for all three example cases when comparing the desired (set) vs. actual (experimental) concentrations (**Figure 7B**). Additionally, the modularity and programmability of the system allows researchers to expose their cultures to any number of different solute concentrations and flow-rate profiles by connecting multiple PiFlow systems in parallel.

**Figure 7:**
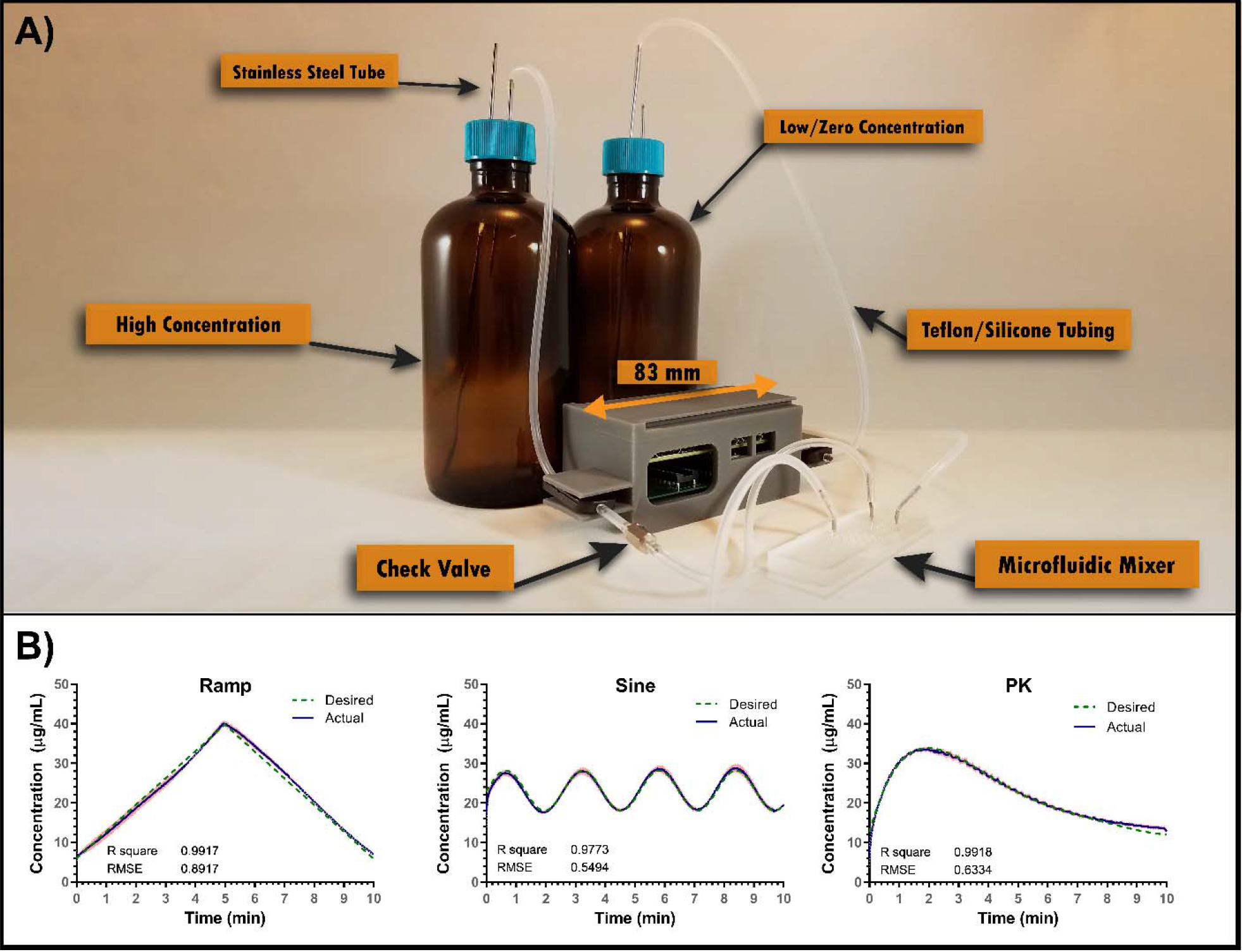
Generating dynamic concentration profiles using PiFlow. **A)** Our typical setup for generating a concentration profile for a single solute. Each pump is connected to a separate reservoir containing the lower and upper limits of the required concentration profile. The flow-rate ratio is then modulated to create the concentration profile at the output of the microfluidic mixer. A unidirectional valve is added to prohibit backflow and make the setup more robust against back pressures. Both silicone and Teflon tubing can be used for the connections. Additionally, thermoplastic-based commercially available mixers can be used in place of our PDMS mixer. **B)** Examples of the experimental concentration profiles generated using fluorescein. The actual mean concentration is represented in blue while the red bands represent the standard deviation. The green dotted line represents the desired (i.e. set) concentration profile. N=3.

### Current Limitations

PiFlow offers a variety of features and capabilities that are desirable to researchers utilizing many kinds of fluidic control. There are several limitations, however, that could be addressed with different designs in the future; 1) Depending on the geometry of the microfluidic device PiFlow is connected to, the flow rates might prove to be too high for certain applications especially those involving very small channels and shear-sensitive cells. While PiFlow can achieve flow rates as low as 1 µL/min, the minimum reliable flow rate highly depends on the setup, including the overall resistance, back pressure and fluid being pumped. Through extended use, we found that the lowest consistently reliable flow rate that can be used with our fully-assembled setup was around 50 µL/min. 2) The current setup uses silicone tubing and a PDMS-based mixer, and while these can be used for a variety of applications, they should not be used in experiments that involve hormones or drugs, as silicone and PDMS tend to be highly absorptive[59–61]. An easy solution would be to use Teflon tubing and a commercially available mixer made from polystyrene or other minimally absorbent plastics although this would significantly increase the cost of operation as commercial mixers cost $70+ per mixer. 3) While PiFlow does have a small footprint, it does not eliminate the need for tubing to connect to the cell-culture devices. Which means as the number of devices increases, so does the number of tubing which can easily become tedious when dealing with a dozen or so devices. 4) While we show the system’s ability to module flow rates on a second-minute time scale, further characterizing by the end user might be required for creating higher frequency components. We have determined that new flow rate values can be sent reliably to the pump at 100 ms intervals, but we have not fully assessed the response time of the system.

### Conclusions

We presented a low-cost (<$350 USD), modular, scalable, fully-programmable and easy to assemble pumping platform that can be used for a variety of fluidic applications, providing all the necessary design and control files to fully implement under an open-source license. We believe our open-source PiFlow system will be valuable to both biological and non-biological researchers, and we look forward to seeing how it will be improved in collaboration with the scientific community.

## Acknowledgements

The authors would like to thank the wider microfluidics community and various researchers that provided both direct and indirect input for the design specifications of PiFlow. This work was funded under the Defense Advanced Projects Agency (DARPA) Microphysiological Systems Program (W911NF-12-2-0039).

